# Application of a Machine Learning Approach Towards the Targeted Identification of Phage Depolymerases

**DOI:** 10.1101/2023.02.28.530424

**Authors:** Damian J. Magill, Timofey A. Skvortsov

## Abstract

Biofilm production plays a clinically significant role in the pathogenicity of many bacteria, limiting our ability to apply antimicrobial agents and contributing in particular to the pathogenesis of chronic infections. Bacteriophage depolymerases, leveraged by these viruses to circumvent biofilm mediated resistance, represent a potentially powerful weapon in the fight against antibiotic resistant bacteria. Such enzymes are able to degrade the extracellular matrix that is integral to the formation of all biofilms and as such would allow complementary therapies or disinfection procedures to be successfully applied. In this manuscript, we describe the development and application of a machine learning based approach towards the identification of phage depolymerases. We demonstrate that on the basis of a relatively limited number of experimentally proven enzymes and using an amino acid derived feature vector that the development of a powerful model with an accuracy on the order of 90% is possible, showing the value of such approaches in the discovery of novel therapeutic agents.

## Background

Biofilms are the most common form of bacterial lifestyle in nature (1). Biofilm formation by pathogenic bacteria allows for the establishment of a multicellular consortium of clinical significance due to the role such communities play in the persistence of bacterial infection and their resistance to various modes of treatment and disinfection. Indeed, such assemblages confer antimicrobial resistance on multiple levels including limiting the penetrability of antimicrobial compounds, the presence of metabolically inactive persister cells exhibiting intrinsic resistance, and the internal structure of such communities providing an optimal environment facilitating horizontal gene transfer (HGT) of resistance determinants (2). A critical component for the establishment of biofilms and a significant contributor to the resistant phenotype they exhibit is the production of a matrix embedding the biofilm cells consisting of various polymeric compounds, including proteins, extracellular DNA, and polysaccharides. The latter can be broadly categorised as lipopolysaccharides (LPS), which are integral components of cell walls of Gram-negative bacteria, capsular polysaccharides (CPS), loosely associated with bacterial surface, and exopolysaccharides (EPS), released by bacteria into the surrounding environment (3). The ability to remove such polymeric barriers in order to expose the underlying community of cells is a desirable one from the practical point of view, be it for the purposes of surface disinfection, de-fouling, or to improve the biocidal effects of antibiotic treatment.

Barrier properties of bacterial biofilms also pose a problem for bacterial viruses (bacteriophages) whose diffusion and ability to infect host cells is reduced within biofilms (4). Targeted degradation of biofilm polysaccharides is a feature of many bacteriophages (phages) which increases the probability of successful infection; this is the result of enzymatic activity of a class of phage-encoded enzymes called depolymerases (DP). The majority of DPs are phage-associated enzymes and belong to lyase and hydrolase classes, with the former constituting a large majority of the well characterised and experimentally validated DPs (5 - 7). Given the global antibiotic crisis we now face, there is a resurgence of interest in both phage and phage-derived therapeutic agents as alternatives. Several recently published reviews describe the structural and functional characteristics of phage DPs and outline their potential applications as biotechnological tools and therapeutic agents (8 - 11).

The therapeutic potential of phage DPs was recognised more than 60 years ago (12). Phage DPs are of particular interest due to their potential use in combinatorial therapies with antibiotics or other antimicrobial agents and in the removal of biofilms from medical devices most notably catheters (13; 14). Moreover, as the depolymerases do not kill bacteria, it is posited that they could be employed on their own as anti-virulence agents, decreasing bacterial fitness and facilitiating the clearance of the bacteria by the human immune system (10). Therefore, any approach that enhances our ability to identify novel DPs is of great value, especially since it is not always trivial to attribute depolymerase activity to a specific gene. As the polysaccharides produced by even closely related bacterial species may have subtle but significant structural differences, phage DPs acting on them also demonstrate high variability, to the point that the depolymerase domains will sometimes be among the only genomic DNA fragments showing no conservation between phages of the same species (14). Although the majority of known DPs are parts of phage receptor-binding proteins (RBPs) such as tail spikes and thus have conserved N-terminal domains responsible for virion attachment, some depolymerases can be encoded as truncated RBPs (presumably acting as diffusible DPs), further complicating their reliable prediction (15).

Machine learning based approaches are proving to be an extremely valuable avenue in all realms of science and this is no less true of phage biology whereby success has been demonstrated through the application of such techniques towards the identification of phage structural proteins (16), host-phage pairs (17), RBPs (18) and lifecycle (19) amongst others (20). Recently published papers expand this list to include endolysins (21) and depolymerases (22). Nevertheless, the ultimate success or failure of machine learning algorithms depends on many factors, including but not limited to the size and composition of training sets, the algorithm used for the problem at hand, and the careful construction of a vector capturing adequately discriminant features (23). Therefore, more ML solutions are needed to expand the computational phage characterisation toolkit and allow for a series of complementary approaches to be available.

In this manuscript we describe the development and application of a machine learning approach towards the identification of phage DPs, highlighting that such models should form an integral part of our toolkit enabling the discovery of novel therapeutics. We demonstrate that even a relatively small training set is sufficient to produce a highly generalizable machine learning model capable of accurately predicting DPs in a multitude of phages infecting vastly different bacteria. Indeed, an accuracy of 90% was attained on the test data set and a similar result for genome context predictions that detected the DP within the top 10 predictions.

## Methods

### Data Set Preparation

In order to establish a database of DP sequences that would ultimately fuel our model, we focussed our attention on publications within which depolymerase activity had been experimentally demonstrated. A comprehensive literature search was conducted and a database established consisting of 50 depolymerase sequences. Table S1 presents an overview of this sequence database including the phages and references from which they were found. 28 of the sequences exclusively state CPS as an enzymatic target, 20 target EPS, and the remaining two target LPS and a combination of targets. The vast majority of sequences were *Podoviridae* derived and the database concerned phages infecting Gram negative bacteria. The size range of sequences varied from 150 amino acids to 1267 amino acids in length.

To complete this dataset, we required 50 sequences that would serve as the negative non-depolymerase set and thus provide a 1:1 positive to negative sequence set. To do this, we randomly extracted 50 sequences from a soil metagenome (SRR15048733) that were sampled across the size distribution of sequences so as to avoid the introduction of sequence size biases. BLAST searches were conducted with these sequences against the positive depolymerases to ensure the absence of homology followed by HHPred analysis to confirm the absence of domains known to be associated with depolymerase activity.

To highlight the dissimilarity in the dataset, we calculated pairwise similarity scores across the entire dataset and represented this as a heatmap (figure 1.).

**Figure 1.**
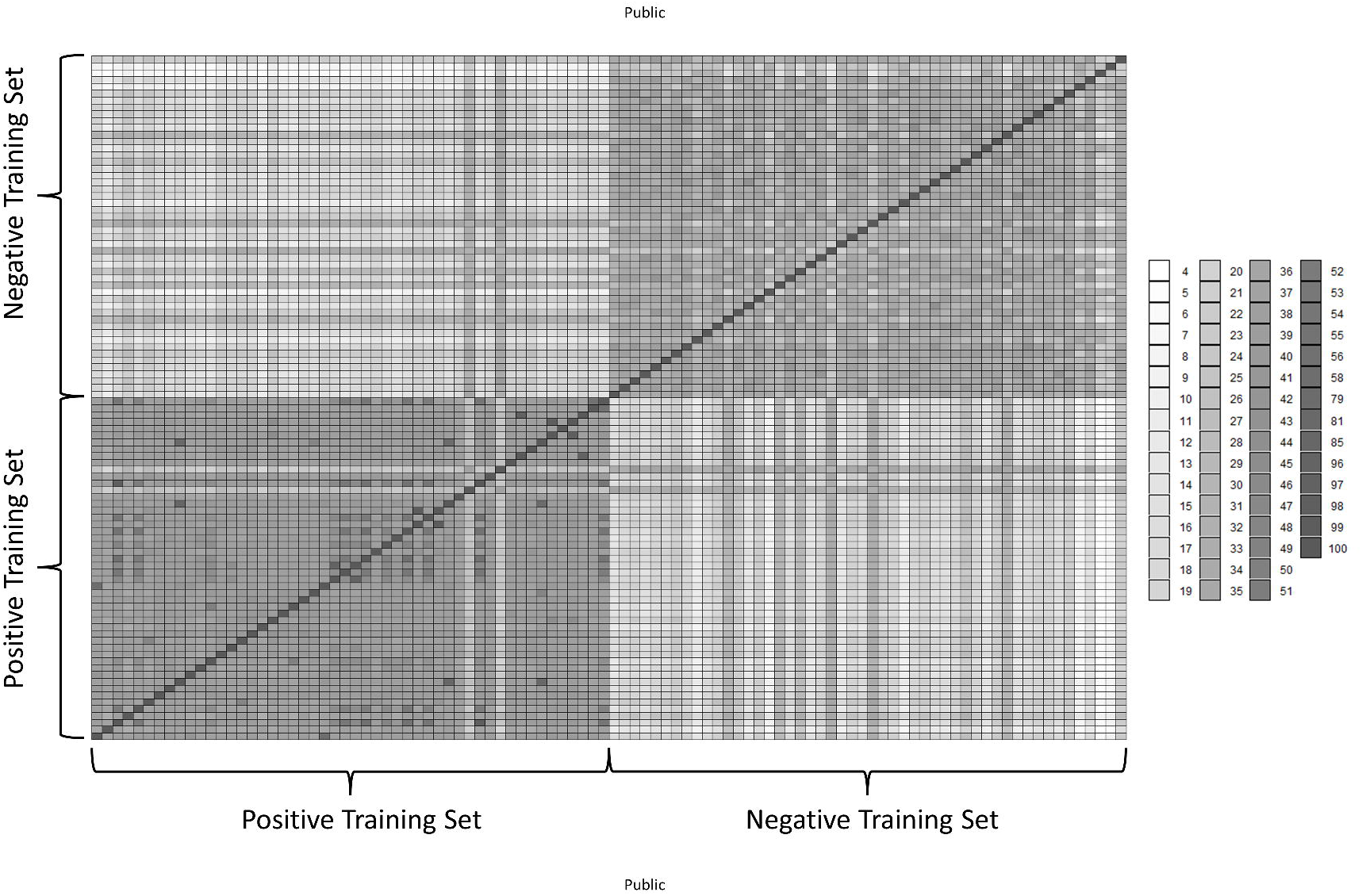
Heatmap of pairwise similarity scores calculated for the training dataset. Grayscale colours correspond to percentage identity as provided in the associated legend. The negative and positive components of the dataset are highlighted with braces and associated labels. As highlighted by the scale of the legend, the global identities of the matrix are rather low, showing a high level of dissimilarity between the sequences.

### Feature Extraction and Selection

A diverse range of features were generated which were derived solely from the amino acid sequences. Eleven of these features were directly calculated using the ProteinAnalysis feature from the BioPython (version 1.73) ProtParam module (24). These were the MW, aromaticity, predicted instability and isoelectric point, GRAVY score, predicted secondary structure (sequence proportion engaged in helices, strands, and turns), extinction coefficients (ox/red), and a combined flexibility score. Beyond this the relative abundance of each amino acid and the total sequence length were also taken into account. As a final set of features, we considered dipeptides and tripeptides as a function of conserved physicochemical properties. Seven groups were established consisting of amino acids with a hydrocarbon R group, those with an uncharged aromatic side chain, sulphur containing, positively charged, negatively charged, polar uncharged, and proline. According to this schema, the dipeptides AE and LD were considered as both belonging to group 15. Whilst allowing us to incorporate dipeptide and tripeptide properties into the model, this also reduced the overall feature set compared to using all possible combinations of amino acids. This was carried out using in-house scripts.

### Model Selection, Training, and Evaluation

With respect to the appropriate choice of machine learning algorithm, we decided to test both support vector machine (SVM) and random forest (RF) approaches (25; 26). This was due to the fact that our data set constituted a small number of samples exhibiting a high feature space. In both cases, we leveraged a grid search in order to assess the hyperparameter space and find the best model configuration for both algorithms. This was conducted using the scikit-learn library (version 0.23.2) (27). We opted for a 5-fold cross validation using and 80/20 split of the dataset.

To evaluate model performance, we particularly focussed on the overall accuracy and recall on the cross-validations defined as follows:

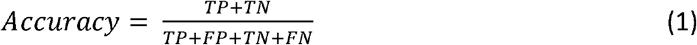

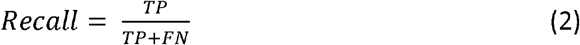

Where TP FP, TN, and FN correspond to true positive, false positive, true negative, and false negative respectively regarding the classification performed on the test data. All scores reported are the average obtained following the cross-validation.

With respect to the hyperparameters tuned, for SVM both linear and RBF kernels were evaluated along with cost and gamma functions when applicable. For RF, differing numbers of estimators were evaluated using a step size of 100 along with total tree depth, and the minimum samples supporting a branch and split of the tree. In addition to this, we also integrated a two-degree polynomial feature transformation, min/max scaling, and applied entropy-based impunity.

Once optimal parameters were determined for the model following evaluation, the final version was created incorporating the entirety of the training set.

### Software Package Depolymerase Predict

Both the source code and a standalone ready to use version of the application are available as detailed in the “Availability of data and materials” section. A simple user-friendly GUI has been developed through which users can step-by-step upload their sequences, generate the feature vector, and carry out predictions and view the output.

## Results and Discussion

### Feature Generation

The application of the feature generation script was carried out on the 100 amino acid sequence input data set. This resulted in the construction of a feature vector with 424 descriptors for each of the sequences. An additional column was added to distinguish the depolymerases from the negative cases. This entire training set is presented in Table S2. and can be used directly in the reproduction of our analysis with the parameters outlined below.

### Model Evaluation and Final Selection

The SVM approach was initially applied to the dataset with no hyperparameter tuning, with the application of a linear kernel. This resulted in a model exhibiting an overall accuracy score of 0.70 across all folds. As presented in the normalised confusion matrix in figure 2. this model performed extremely poorly with respect to true and false positives but handled negative cases well. Indeed, hyperparameter tuning did nothing to resolve this problem. The overall accuracy remained unchanged, but the model improved in its ability to correctly identify non-depolymerase sequences with 100% success rate. This was at the cost of decreased performance on positive cases with only 45% of true depolymerases being correctly identified as such.

**Figure 2.**
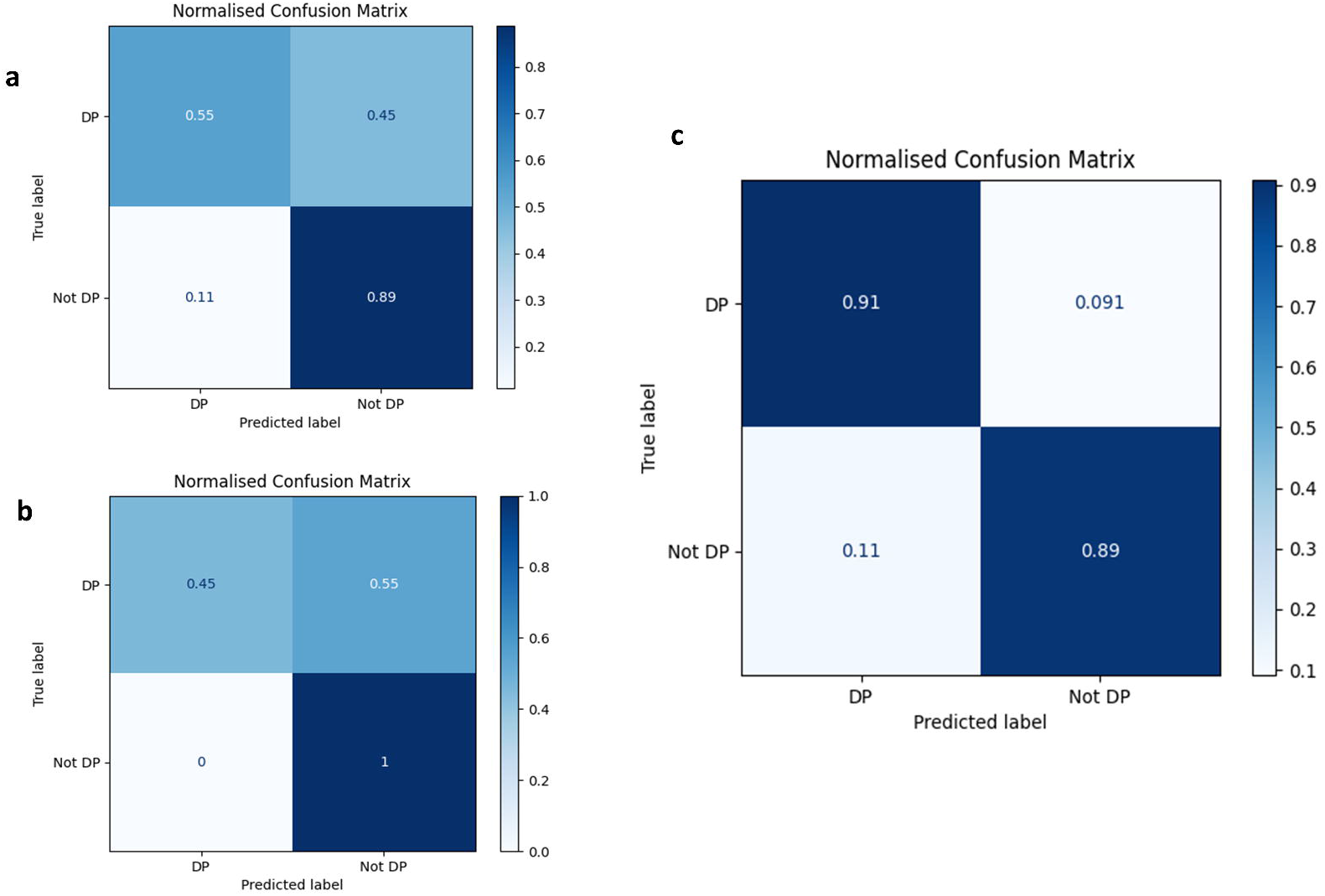
Normalised confusion matrices summarising model performance on test data. Matrices give the proportion of depolymerase (DP) and non-depolymerase (Not DP) that are correctly identified by the model, corresponding thus to the true/false positive and true/false negative proportions. Matrices are shown for non-optimised SVM (a), optimised SVM (b), and optimised RF (c) models.

Subsequent application of an RF approach yielded more promising results. This is an ensemble machine learning method that leverages multiple decision trees in order to reduce variance and provide better model generalization. It performs especially well with small sample sets and large feature spaces and so it was expected it may be the best approach to this problem. Application of a tuned RF model indeed showed a much higher level of performance (figure 2). An overall accuracy score of 0.90 was obtained across all folds with similar performance observed with respect to the correct classification of positive and negative cases. It was found that for this case, the following parameters provided optimal performance of the model: use of 1500 estimators with automatic definition of maximum features to be used by each tree. A maximum depth of 30 was applied with a minimum sample support of 3 required for each leaf. Each tree split was evaluated using the entropy-based criterion. The pipeline also integrated a two-degree polynomial feature transformation along with application of min/max scaling.

### Application of Model Towards Depolymerase Identification

In order to further assess the performance of our model, we decided to apply it within the context of whole phage genomes to see whether it could correctly identify depolymerases amongst the other genes. Due to the fact that our model leverages experimentally active depolymerases and we have thus exhausted this option, we were limited to performing this test on computationally predicted enzymes. The first such case was *Pseudomonas* phage pf16; a phage previously characterised by our group (28). Depolymerase activity was previously observed in this phage and extensive computational analysis identified gp215 as the likely candidate with probable pectate lyase activity. We proceeded to analyse pf16 gene products using our model and ranked the probabilities accordingly. These results are presented in figure 3. We immediately observed that the predicted depolymerase was ranked 4^th^ by our model in the context of the whole genome. This in itself is a reasonable result however, further analysis of the higher ranked candidates revealed that they possess domains not unrelated to what is observed in depolymerases including endosialidase and VrlC-like domains, the latter speculated to have sialidase activity (29).

**Figure 3.**
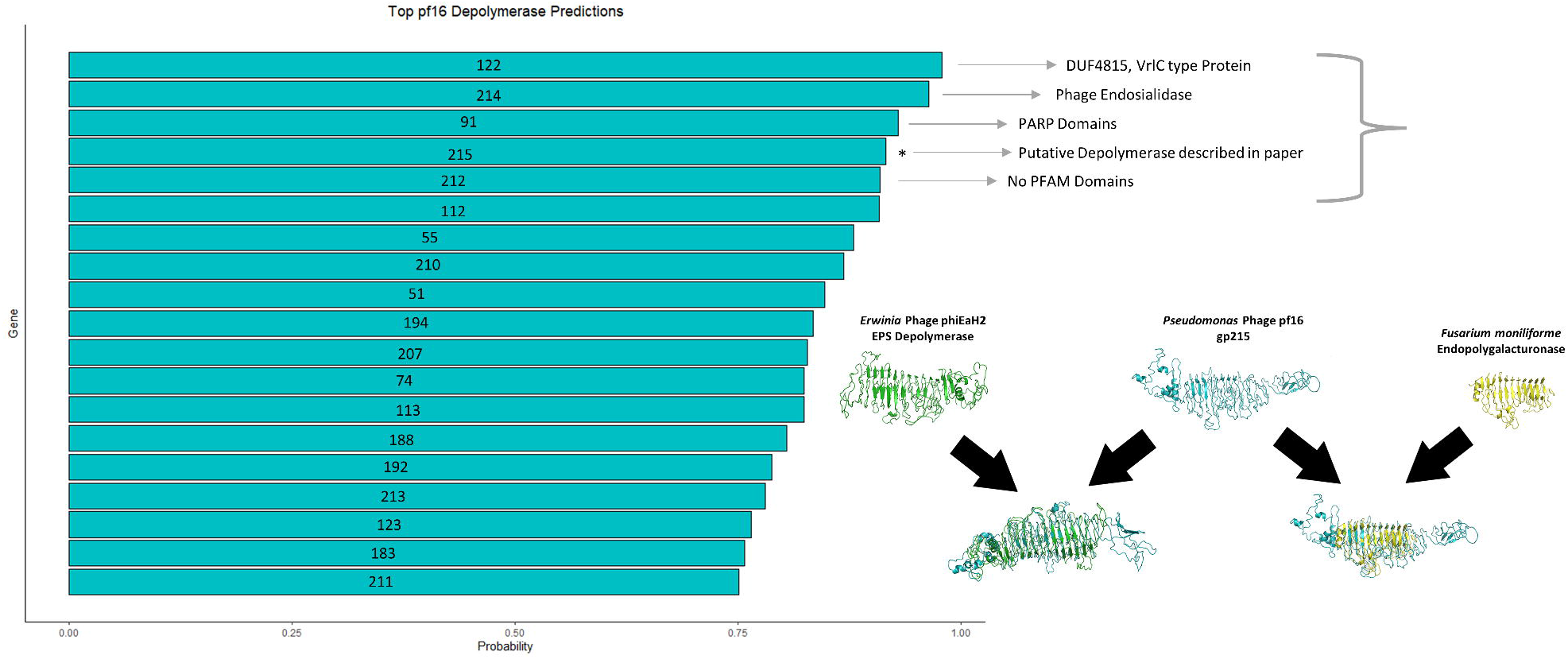
Top Predictions of *Pseudomonas* phage pf16 depolymerases. The graph highlights that probability reported by the model of the gene product being a depolymerase. Gene products are labelled accordingly. The putative depolymerase previously reported is highlighted on the graph and the modelling of this protein shown with respect to a known EPS depolymerase and endopolygalacturonase as reported in *Magill* et al. (2017).

We further tested the performance of our model in the context of whole genomes by directing our attention towards computationally predicted depolymerases described by Pires *et al*. (11). This provided a good opportunity to test the generalizability of our model as the sequences described in this paper exhibit significant diversity in terms of the domains present and nature of the hosts infected by the phages. We downloaded the genomes of the associated phages, removing some for which the records no longer exist. This resulted in 155 genomes on which we applied our model. Predictions were performed, the probabilities ranked, and the position of the putative depolymerase identified. Table S3 presents all of the genomes, the depolymerases and the associated ranking provided by our model. Across all sequences the depolymerase featured as the first prediction 40.6% of the time. This increased to 69.7% and 78.1% for top 3 and top 5 predictions respectively. When considering top 10 and top 20 this grows to a large majority with 87.1% and 94.8%. Most poorly predicted sequences were those containing domains that did not feature in our model, especially DUF867. When we look closer at the distribution of these results we observe a good level of model generalizability in a number of aspects (Figure 4). Despite being fuelled by depolymerases in phages infecting Gram-negative bacteria, the model performs equally well for phages infecting both types. This fact also holds when considering the family of phage and the genus of the host. This implies that the model is leveraging features that are common to a large majority of known depolymerase enzymes.

**Figure 4.**
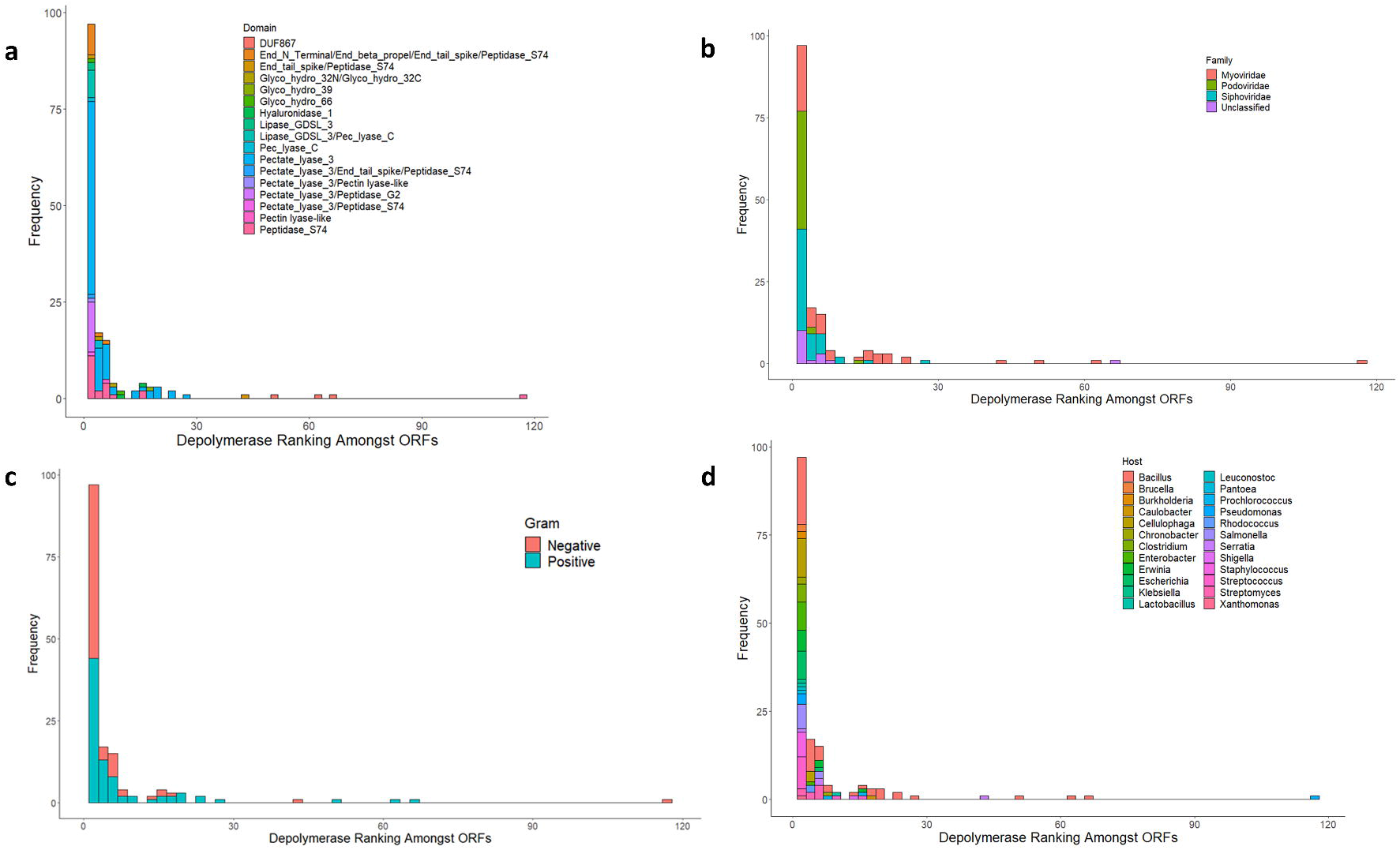
Graphs showing ranking of depolymerases predicted by the model. Rankings performed on depolymerase predictions from genomes described by Pires *et al*. (2016). Rankings are coloured by depolymerase domains (a), family of the phage described (b), whether the host is Gram-positive or negative (c), and by the host genus (d).

## Conclusion

Bacteriophage depolymerases offer a host of promising clinical and biotechnological applications, including the synergistic treatment of infections via biofilm removal. There is however, a need for rapid and accurate identification of such enzymes. In this work we have described the development and application of a machine learning approach that allows for depolymerase prediction with an overall accuracy of 90% using a sequence-derived feature vector. We demonstrated that this model was generalizable to depolymerases from a variety of phages, robustly predicting them in the context of the genomes across several hosts and enzyme classes. This highlights the power that such approaches can offer in the identification of industrially and/or clinically useful enzymes.

## Declarations

### Ethics approval and consent to participate

Not applicable

### Consent for publication

Not applicable

### Availability of data and materials

The source code for the application can be found via the following URL: https://github.com/DamianJM/Depolymerase-Predict.git

In addition, a standalone version of the application is available with all dependencies and training set compiled within at the following address: https://sourceforge.net/projects/depolymerase-predict

The training dataset has been provided as part of the supplementary data which allows for our work to be reproduced. Depolymerases used in the development of the model are detailed in Supplementary table 1, with all accession numbers and associated literature references provided.

### Authors’ contributions

DM developed the software; DM and TS analysed the data; DM and TS wrote the manuscript and approved the final version along with figures.

## Acknowledgements

Not applicable

