## Supplementary material for "Application of a Machine Learning Approach Towards the Targeted Identification of Phage Depolymerases": Table S1.

| **Phage** | **Bacterial Host** | **Family** | **Genus** | **Depolymerase target** | **Accession no.** | **DNA sequence** | **Gene Location** | **Reference** |
| --- | --- | --- | --- | --- | --- | --- | --- | --- |
| IME200 | Acinetobacter baumannii | Autographivirinae | Fri1virus | EPS | ALJ97635 | 35572..37653 |  | (Liu *et al.*, 2019) |
| GH-K3 | Klebsiella pneumoniae | Siphoviridae |  | CPS | AYP28214 | 21164..24967 | GHK3_33 | (Cai *et al.*, 2019) |
| SH-KP152226 | Klebsiella pneumoniae | Podoviridae Autographivirinae |  | EPS | QDF14644 | 33566..35947 | SHKP152226_42 | (Wu *et al.*, 2019) |
| IME180 | Pseudomonas aeruginosa | Podoviridae Caudovirales |  | EPS | ATG86239 | 861..2255 |  | (Mi *et al.*, 2019) |
| KP32 | Klebsiella pneumoniae | Podoviridae |  | CPS K3 | YP_003347555 | 33420..36029 | KP-KP32_gp37 | (Majkowska-Skrobek *et al.*, 2018) |
| KP32 | Klebsiella pneumoniae | Podoviridae |  | CPS K21 | YP_003347556 | 36040..37770 | KP-KP32_gp38 | (Majkowska-Skrobek *et al.*, 2018) |
| vB_EcoM_ECOO78 | Escherichia coli | Myoviridae |  | EPS | ARM70447 | 27628..29871 | vBEcoMECOO78_42 | (Guo *et al.*, 2017) |
| KP36 | Klebsiella pneumoniae | Siphoviridae | KP36likevirus | CPS | YP_009226011 | 29509..32160 | AXI63_gp50 | (Majkowska-Skrobek *et al.*, 2016) |
| IME285 | Acinetobacter baumannii | Myoviridae |  | CPS | AYP68900 | 22578..24776 |  | (Wang *et al.*, 2020) |
| Petty | Acinetobacter nosocomialis and Acinetobacter baumannii | Podoviridae |  | EPS | AGY48011 | 33863..36619 | Gp39 | (Hernandez-Morales *et al.*, 2018) |
| PHB02 | Pasteurella multocida |  |  | CPS | ARV77571 | 4014..6920 | Gp17 | (Chen *et al.*, 2018) |
| kpssk3 | Carbapenem-resistant Klebsiella pneumoniae | Podoviridae  Autographivirinae | Teseptimavirus | EPS | AZF88843 | 33770..36823 | kpssk3_038 | (Shi *et al.*, 2020) |
| ΦK64-1 | Multi-host  Klebsiella pneumoniae | Myoviridae | Klebsiella | EPS | BAQ02805 | 90154..91941 |  | (Pan *et al.*, 2017) |
| ΦK64-1 | Multi-host  Klebsiella pneumoniae | Myoviridae | Klebsiella | EPS | BAQ02835 | 321593..323803 |  | (Pan *et al.*, 2017) |
| ΦK64-1 | Multi-host  Klebsiella pneumoniae | Myoviridae | Klebsiella | EPS | BAQ02836 | 323855..325810 |  | (Pan *et al.*, 2017) |
| ΦK64-1 | Multi-host  Klebsiella pneumoniae | Myoviridae | Klebsiella | EPS | BAQ02837 | 325887..327995 |  | (Pan *et al.*, 2017) |
| ΦK64-1 | Multi-host  Klebsiella pneumoniae | Myoviridae | Klebsiella | EPS | BAQ02838 | 328067..331648 |  | (Pan *et al.*, 2017) |
| ΦK64-1 | Multi-host  Klebsiella pneumoniae | Myoviridae | Klebsiella | EPS | BAQ02839 | 331729..333483 |  | (Pan *et al.*, 2017) |
| ΦK64-1 | Multi-host  Klebsiella pneumoniae | Myoviridae | Klebsiella | EPS | BAQ02841 | 335902..338568 |  | (Pan *et al.*, 2017) |
| ΦK64-1 | Multi-host  Klebsiella pneumoniae | Myoviridae | Klebsiella | EPS | BAQ02842 | 338917..341907 |  | (Pan *et al.*, 2017) |
| ΦK64-1 | Multi-host  Klebsiella pneumoniae | Myoviridae | Klebsiella | EPS | BAQ02844 | 344312..346471 |  | (Pan *et al.*, 2017) |
| B9 | Acinetobacter baumannii | Myoviridae | R2096virus | EPS | AWD93192 | 40985..43576 | AB9_069 | (Oliveira *et al.*, 2018) |
| ΦAB6 | Acinetobacter baumannii | Autographiviridae | Friunavirus | EPS | ALA12264 | 34281..36380 | phiAB6_gp40 | (Lai *et al.*, 2016) |
| IME321 | Klebsiella pneumoniae | Podoviridae | Kp32 virus | CPS | AXE28435 | 33394..35856 |  | (Wang *et al.*, 2019) |
| K30 | Escherichia coli | Autographiviridae | Przondovirus | CPS | YP_004678762 | 32920..35631 | EnPhK30_gp41 | (Lin *et al.*, 2017) |
| KN1-1 | Klebsiella pneumoniae | Autographiviridae | Przondovirus | CPS | BBF66844 | 34320..36782 | KN1dep | (Pan *et al.*, 2019) |
| KN3-1 | Klebsiella pneumoniae | Autographiviridae | Przondovirus | CPS | BBF66867 | 33247-35622 | KN3dep | (Pan *et al.*, 2019) |
| KN3-1 | Klebsiella pneumoniae | Autographiviridae | Przondovirus | CPS | BBF66868 | 35635-37668 | K56dep | (Pan *et al.*, 2019) |
| KN4-1 | Klebsiella pneumoniae | Autographiviridae | Przondovirus | CPS | BBF66888 | 34859-37408 | KN4dep | (Pan *et al.*, 2019) |
| PHB19 | Shiga toxin-producing Escherichia coli | Autographivirinae |  | CPS, LPS, EPS | QHI00738 | 4001..7078 |  | (Yibao *et al.*, 2020) |
| K5-2 | Klebsiella pneumoniae | Autographiviridae | Przondovirus | CPS | APZ82804 | 33140..35518  k52_037 |  | (Hsieh *et al.*, 2017) |
| K5-2 | Klebsiella pneumoniae | Autographiviridae | Przondovirus | CPS | APZ82805 | 35529..37586 | k52_038 | (Hsieh *et al.*, 2017) |
| K5-4 | Klebsiella pneumoniae | Autographiviridae | Przondovirus | CPS | APZ82847 | 32318..34567 | k54_037 | (Hsieh *et al.*, 2017) |
| K5-4 | Klebsiella pneumoniae | Autographiviridae | Przondovirus | CPS | APZ82848 | 34578..36632 | k54_038 | (Hsieh *et al.*, 2017) |
| AM24 | Acinetobacter  baumannii | Myoviridae |  | CPS | APD20249 | 19770..22316 | AM24_050 | (Popova *et al.*, 2019) |
| S2 | Erwinia amylovora |  | SP6virus | EPS | AUV57247 | 42745..44718 |  | (Knecht *et al.*, 2018) |
| Bue1 | Erwinia amylovora | Ackermannviridae |  | EPS | AVO22848 | 5408..5935 |  | (Knecht *et al.*, 2018) |
| IME205 | Klebsiella pneumoniae | Autographiviridae | Przondovirus | CPS | ALT58497 | 33550..35931 | Orf42 | (Liu *et al.*, 2020) |
| IME205 | Klebsiella pneumoniae | Autographiviridae | Przondovirus | CPS | ALT58498 | 35950..37875 |  | (Liu *et al.*, 2020) |
| phiAp1 | Ralstonia spp | Podoviridae | Phikmvvirus | EPS | APU03184 | 29924..30376 | phiAp1_43 | (da Silva Xavier *et al.*, 2018) |
| kpv71 | Klebsiella pneumoniae | Podoviridae | Kp34virus | CPS | AMQ66478 | 40466..42421 | kpv71_52 | (Solovieva *et al.*, 2018) |
| kpv74 | Klebsiella pneumoniae | Podoviridae | Kp34virus | CPS | APZ82768 | 42255..43988 | kpv74_56 | (Solovieva *et al.*, 2018) |
| PP35 | Dickeya solani | Ackermannviridae | Limestonevirus | LPS | ATW62160 | 120600..122246 | orf156 | (Kabanova *et al.*, 2019) |
| BS46 | Acinetobacter  Baumannii | Myoviridae |  | CPS | QEP53229 | 33168..35645 | BS46_gp47 | (Popova *et al.*, 2020) |
| πVLC5 | Klebsiella pneumoniae | Podoviridae | Drulisvirus | CPS | QIW86419 | 36103..38478 | VLC5_49 | (Domingo-Calap *et al.*, 2020) |
| πVLC5 | Klebsiella pneumoniae | Podoviridae | Drulisvirus | CPS | QIW86428 | 42631..44634 | VLC5_58 | (Domingo-Calap *et al.*, 2020) |
| πVLC6 | Klebsiella pneumoniae | Podoviridae | Drulisvirus | CPS | QJI52623 | 35765..38140 | VLC6_51 | (Domingo-Calap *et al.*, 2020) |
| πVLC6 | Klebsiella pneumoniae | Podoviridae | Drulisvirus | CPS | QJI52632 | 42292..44025 | VLC6_58 | (Domingo-Calap *et al.*, 2020) |
| KpV79 | Klebsiella pneumoniae | Autographiviridae |  | CPS | ATI16495 | 26936..29101 | kpv79_42 | (Volozhantsev *et al.*, 2020) |
| kpv767 | Klebsiella pneumoniae | Autographiviridae |  | CPS | AOZ65519 | 34114..36645 | kpv767_46 | (Volozhantsev *et al.*, 2020) |
